## Supplementary Information for "Modeling and mechanical perturbations reveal how spatially regulated anchorage gives rise to spatially distinct mechanics across the mammalian spindle"

#### Table of Contents

**A Euler-Bernoulli formalism and generation of native k-fiber shapes** **S2**

A.1 Euler-Bernoulli equation and the small angle approximation . . . . . S2

A.2 Analytical solutions for native k-fiber profiles . . . . . S2

A.3 Justification of the small angle approximation . . . . . S4

**B Justification of treating the microneedle force as a point force** **S6**

**C Numerical approach for solving the Euler-Bernoulli equation** **S7**

**D Model fitting procedure** **S8**

**E Modeling of PRC1 binding states** **S9**

### A Euler-Bernoulli formalism and generation of native k-fiber shapes

In this section, we use the Euler-Bernoulli formalism to calculate the shapes of native k-fiber profiles (see Fig. 2c of the main text for example profiles). In our analysis, we consider k-fibers at mechanical equilibrium and assume that their shapes are generated by forces and moments acting at their end-points. All calculations are performed in the reference frame where the pole and the kinetochore lie along the $x$-axis. The calculations and results of this section are related to Fig. 2 of the main text.

#### A.1 Euler-Bernoulli equation and the small angle approximation

The Euler-Bernoulli beam theory relates the local curvature $\kappa(x)$ to the bending moment $M(x)$ via

$$\kappa(x) = -\frac{M(x)}{EI}, \tag{1}$$

where $EI$ is the flexural rigidity of the beam, with $E$ being the Young's modulus, and $I$ being the areal moment of inertia. The general expression for the curvature written in terms of Cartesian coordinates is given by

$$\kappa(x) = \frac{y''(x)}{\left(1 + y'(x)^2\right)^{3/2}}. \tag{2}$$

When substituted into Eq. 1, this expression for $\kappa(x)$ results in a nonlinear equation for the k-fiber profile $y(x)$, making it challenging to obtain an analytical solution and extract intuition from it. We therefore begin our calculations by making the so-called 'small angle approximation' in order to write $\kappa(x) \approx y''(x)$. This approximation applies to the native k-fibers shapes, which, upon aligning them along the $x$-axis, appear flat and have a small tangential angle at every position of the profile (i.e., $|y'(x)| \ll 1$). This leads to a simpler and analytically tractable form for the Euler-Bernoulli equation, namely,

$$y''(x) = -\frac{M(x)}{EI}. \tag{3}$$

Solving this simpler equation will let us gain insights into how the different model components that define $M(x)$ uniquely contribute to k-fiber shape. Later in section A.3, we will demonstrate the validity of applying the 'small angle approximation' for native k-fiber shapes.

#### A.2 Analytical solutions for native k-fiber profiles

In our minimal model, the shape of native k-fibers is generated due to forces and moments acting at the pole and kinetochore ends of the k-fiber (Fig. 2b of the main text). The $x$- and $y$-components of the force at the pole are denoted by $F_x$ and $F_y$, respectively. The bending moment at the pole is denoted by $M_p$, with the counterclockwise direction chosen to be positive. The bending moment at the kinetochore ($M_k$) is generally different from $M_p$.

Eq. 3 is a second-order ordinary differential equation for the k-fiber shape $y(x)$. To obtain $y(x)$, we need to specify two boundary conditions. These condition are

$$y(0) = 0, \tag{4}$$
$$y(L) = 0, \tag{5}$$

where $L$ is the distance between the pole and kinetochore ends of the k-fiber. These conditions require that the k-fiber ends are positioned on the $x$-axis.

The local bending moment $M(x)$ is obtained by writing the moment balance condition for the $[0, x]$ segment of the k-fiber (see Fig. 2b of the main text). Specifically, $M(x)$ needs to balance the torque generated by the force $\vec{F} = (F_x, F_y)$ at the pole and the bending moment $M_p$, i.e.,

$$M(x) = M_p + \underbrace{F_x\, y(x) - F_y\, x}_{\left(\vec{r}\times\vec{F}\right)_z}. \tag{6}$$

Using the above expression, we can relate the bending moment at the kinetochore ($M_k \equiv M(x = L)$) to the moment at the pole ($M_p$). Noting that $y(L) = 0$, we obtain

$$M_k = M_p - F_y L. \tag{7}$$

This indicates that a non-zero end-point force perpendicular to the pole-kinetochore axis will necessarily result in different bending moments (hence, curvatures) at k-fiber ends.

Knowing how the bending moment varies in space (Eq. 6), we now substitute it into the Euler-Bernoulli equation in its 'small angle approximation' form (Eq. 3) and obtain a linear second order ODE for the profile function $y(x)$, namely,

$$y''(x) + \left(\frac{F_x}{EI}\right) y(x) = -\frac{M_p}{EI} + \left(\frac{F_y}{EI}\right) x. \tag{8}$$

We note that the forces and the moment at the pole always appear in a ratio with the flexural rigidity $EI$. We therefore introduce rescaled effective parameters $\tilde{F}_x = F_x/EI$, $\tilde{F}_y = F_y/EI$, and $\tilde{M}_p = M_p/EI$, and rewrite the second order ODE for $y(x)$ as

$$y''(x) + \tilde{F}_x y(x) = -\tilde{M}_p + \tilde{F}_y x. \tag{9}$$

The functional form of the solution for $y(x)$ depends on the signs and values of the different parameters. Therefore, in the following, we consider separate scenarios and discuss the insights that each analytical solution provides.

$\underline{F_x = 0}$. We begin with the special case where the point force along the pole-kinetochore axis is zero. This simplifies the differential equation into

$$y''(x) = -\tilde{M}_p + \tilde{F}_y x. \tag{10}$$

Integrating twice over $x$, we obtain

$$y(x) = C_1 + C_2 x - \frac{\tilde{M}_p x^2}{2} + \frac{\tilde{F}_y x^3}{6}, \tag{11}$$

where $C_1$ and $C_2$ are integration constants. Imposing the boundary conditions (Eq. 4 and Eq. 5), we find these constants to be $C_1 = 0$ and $C_2 = \tilde{M}_p L/2 - \tilde{F}_y L^2/6$. Substituting $C_1$ and $C_2$, and writing the perpendicular force as $\tilde{F}_y = (M_p - M_k)/L$ (from Eq. 7), we obtain the final expression for $y(x)$:

$$y(x) = \tilde{M}_p \underbrace{\frac{x(L-x)}{2}}_{\text{symmetric}} - (\tilde{M}_p - \tilde{M}_k) \underbrace{\frac{x(L^2-x^2)}{6L}}_{\text{asymmetric}}. \tag{12}$$

The first term contributing to the profile is symmetric about the middle position $x = L/2$ and does not change under the transformation $x \to L - x$. The second term, however, is asymmetric and leads to a shift of the profile peak toward the end which has the higher bending moment.

In the limit where there is bending moment at the pole but not at the kinetochore ($\tilde{M}_p > 0$, $\tilde{M}_k = 0$), we can find the peak position of the asymmetric profile by solving for $x$ in the equation $y'(x) = 0$. We obtain $x_{\text{peak}} = \left(1 - \sqrt{3}/3\right) L \approx 0.42\, L$, which means that the peak of the profile is shifted toward the pole side by $\approx 8\%$ of the end-to-end distance $L$. Similarly, when bending is present only at the kinetochore ($\tilde{M}_p = 0$, $\tilde{M}_k > 0$), the profile peaks at $x \approx 0.58\, L$, which is shifted now toward the kinetochore side by the same amount (see Fig. 2c of the main text for demonstrations of these two asymmetric cases).

$\underline{F_x > 0 \text{ and } M_p = M_k = 0}$. Next, we consider another special case where the k-fiber profile is formed by a purely axial force $F_x$ in the absence of bending moments at either end-point ($\tilde{M}_p = \tilde{M}_k = 0$ and hence, $\tilde{F}_y = 0$). The ODE for $y(x)$ in this case simplifies into

$$y''(x) + \tilde{F}_x\, y(x) = 0. \tag{13}$$

The general solution is a linear combination of $\sin(kx)$ and $\cos(kx)$ functions, with the wave number defined as $k = \sqrt{\tilde{F}_x}$. The boundary condition $y(0) = 0$ eliminates the cosine solution. Imposing the second boundary condition, we obtain $\sin(kL) = 0 \Rightarrow k = \pi/L$ (first buckling mode). This suggests that the axial force needs to exactly equal the critical buckling force given by $F_c = \pi^2 EI/L^2$. The sinusoidal buckling profile, as shown in Fig. 2c, is symmetric with respect to $x = L/2$.

$F_x > 0$, $M_p > 0$, and $M_k = 0$. We end our analytical treatment of native k-fiber shapes by considering the more general case where a moment at the pole and axial forces are both present, but there is no moment generation at the kinetochore ($\tilde{M}_k = 0$). This corresponds to the minimal model sufficient to capture the diverse shapes of k-fibers in their native state (see Fig. 2f,g of the main text).

Substituting $\tilde{F}_y = \tilde{M}_p/L$ into Eq. 9, the ODE for the k-fiber profile $y(x)$ for this case becomes

$$y''(x) + \tilde{F}_x y(x) = -\frac{\tilde{M}_p}{L}(L - x). \tag{14}$$

The general solution to the ODE can be written as

$$y(x) = D_1 \sin\big(k(L - x)\big) + D_2 \cos\big(k(L - x)\big) - \frac{\tilde{M}_p}{\tilde{F}_x L}(L - x), \tag{15}$$

where $D_1$ and $D_2$ are integration constants. The argument of sine and cosine functions is written as $(L - x)$ for convenience. Now, the boundary condition at the kinetochore is $y(x = L) = 0$ which indicates that $D_2 = 0$. From a similar boundary condition at the pole ($y(0) = 0$), we obtain $D_1 = (\tilde{M}_p/\tilde{F}_x)/\sin(kL)$. After substitution, the final expression for the profile becomes

$$y(x) = \frac{\tilde{M}_p}{\tilde{F}_x}\left(\frac{\sin(k(L - x))}{\sin(kL)} - \frac{L - x}{L}\right), \tag{16}$$

where, as a reminder, $k = \sqrt{\tilde{F}_x}$. One can show that in the limit where the axial force goes to zero ($k \to 0$), the polynomial solution in Eq. 12 is recovered. Conversely, when the axial force is close to the critical buckling force (achieved when $kL \approx \pi$ or $\tilde{F}_x \approx \pi^2/L^2$), the sine term becomes dominant in the solution and the symmetric sinusoidal profile is recovered (solution of Eq. 13).

To probe the behavior in the intermediate regimes, we tuned the axial force in the $(0, F_c)$ range and numerically found the corresponding moment at the pole ($M_p$) that would yield an identical peak deflection, which we set equal to $y_{\max} = 0.1L$ (our conclusions hold true for any other $y_{\max}$ value that does not violate the small-angle approximation). As shown in Fig. S1a, the larger the axial force becomes, the smaller the corresponding moment at the pole needs to be in order to yield the same amount of k-fiber deformation (measured by $y_{\max}$). Furthermore, as anticipated, increasing the axial force (with a corresponding decrease in the moment at the pole) shifts the position of the peak closer to the center (Fig. S1b), and when the force reaches the critical buckling force, the $x$-position of the peak becomes equal to $L/2$.

Overall, this study shows that the simultaneous tuning of $M_p$ and $F_x$ (equivalently, $F_y$ and $F_x$) according to the rule revealed in Fig. S1a shifts the $x$-position of the k-fiber peak without changing the magnitude of the peak k-fiber deflection ($y_{\max}$).

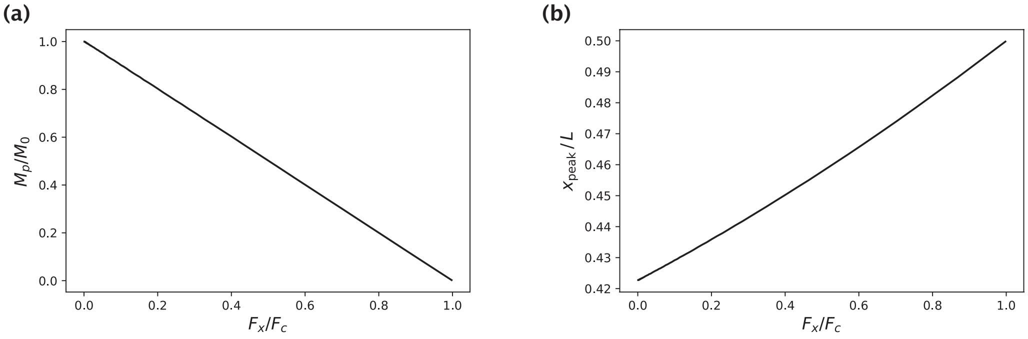


Figure S1: Generation of native k-fiber shapes with a fixed peak deflection $y_{\max}$ for different choices of the axial force $F_x$. (a) Moment at the pole $M_p$ that yields the specified peak deflection $y_{\max}$ as a function of $F_x$. $M_0$ is the pole moment in the absence of an axial force. (b) $x$-position of the profile peak as a function of $F_x$.

#### A.3 Justification of the small angle approximation

Here, we demonstrate the validity of the small angle approximation for modeling native k-fiber shapes by comparing the results of inference under this approximation with the results of the more exact numerical approach detailed in section C.

In our minimal model of native k-fiber shape generation, the two independent parameters are the axial force $F_x$ and the moment at the pole $M_p$. For each parameter, we take the ratio of its inferred value under the approximate and exact methods, and plot the value of this ratio for all native k-fibers considered in our study (Fig. S2a). When plotting, we give transparency to each data point based on how much they contribute to k-fiber shape. If the data point is transparent, then the corresponding parameter ($F_x$ or $M_p$) contributes little to k-fiber shape. We set the weights of shape contribution for axial force and moment at the pole as $w_{F_x} = |F_x|y_{\text{max}}/(|F_x|y_{\text{max}} + M_p)$ and $w_{M_p} = M_p/(|F_x|y_{\text{max}} + M_p)$, respectively, where $|F_x|y_{\text{max}}$ is the largest mechanical moment exerted by the axial force about the origin $(0, 0)$. As can be seen from Fig. S2a, parameters inferred by the two methods are almost always very close to each other when the corresponding parameter has a significant shape contribution, and may differ significantly when the corresponding parameter does not contribute significantly to shape (transparent points). Furthermore, the fitting errors predicted by the two methods are very similar to each other (Fig. S2b). Together, these studies demonstrate the validity of invoking the small angle approximation for studying native k-fiber shapes.

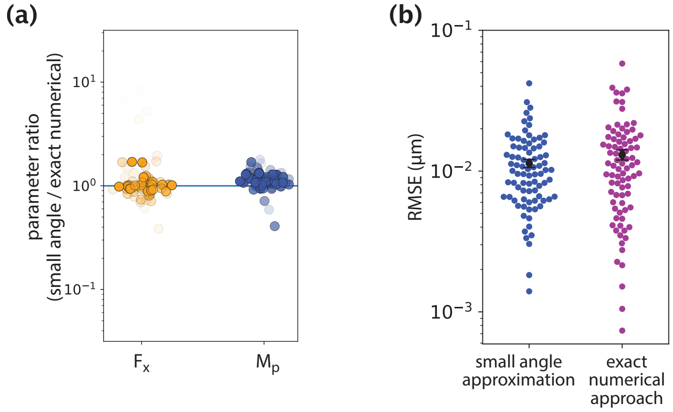


Figure S2: Comparison of inference results from the small angle approximation and exact numerical approach. (a) Ratios of parameters inferred by the two methods. (b) Fitting error comparison between the two methods.

### B Justification of treating the microneedle force as a point force

In this section, we present the results of the studies that justify our treatment of the microneedle force as a point force in models of k-fiber shape generation (related to Figs. 3-5 in the main text). This is in contrast to using a distributed force acting along a finite region of the k-fiber, the size of which is set by the diameter of the microneedle ($\approx 0.5$–$1.5\,\mu$m).

To demonstrate that k-fiber response is not significantly dependent on the distributed force assumption, we first generated k-fiber shapes by applying distributed forces over regions of varying size (0.5–1.5 µm) and overlaid the resulting profiles. As shown in Fig. S3, there is a very close match between the k-fiber profiles generated by the same integrated force distributed along different length scales.

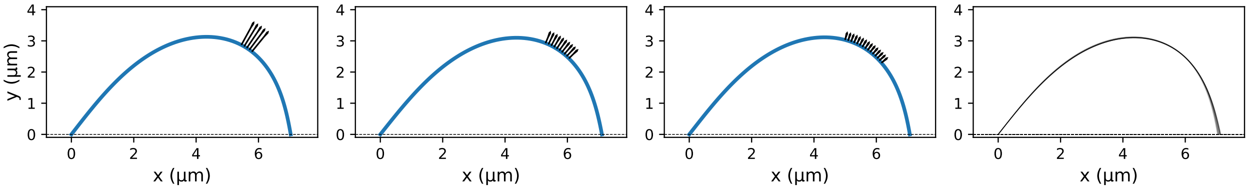


Figure S3: K-fiber profile generation with a distributed microneedle force. (a) Example profiles where the same integrated microneedle force was applied over three different regions of the k-fiber. The black arrows indicate the applied external forces with lower magnitudes for larger regions of distributed force application. Parameters used in profile generation: $\tilde{F}_{\text{ext}} = 0.2\,\mu\text{m}^{-2}$ (integrated force), $L_{\text{contour}} = 10$ µm. (b) Profiles from 10 different distributed force settings overlaid on top of each other. Only minor differences can be observed near the rightmost end.

To further validate our point-force treatment, we inferred a point-force model for the synthetically generated profiles with distributed forces (see sections C and D for details on the inference procedure). In all cases, the effective point force was inferred to be within $\approx 0.02\,\mu$m of the center of the distributed force application region (Fig. S4), with the error between the generated data and inferred model being very low (RMSE $\approx 0.03\,\mu$m). Together, these results justify our treatment of the microneedle force as a point force in our models of k-fiber manipulation.

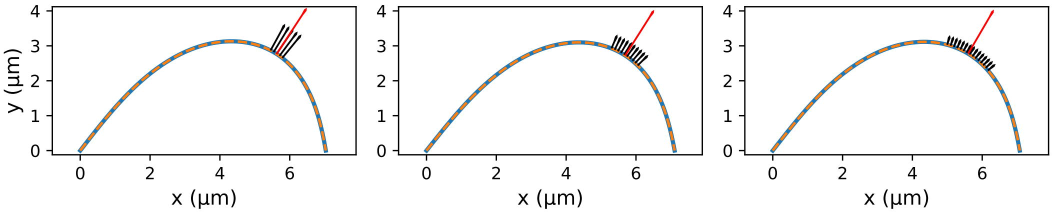


Figure S4: Fits of the point microneedle force model (dashed lines) to synthetic profiles (solid blue lines) which were generated with distributed forces applied over different regions. The red arrow indicates the inferred point microneedle force $F_{\text{ext}}$.

### C Numerical approach for solving the Euler-Bernoulli equation

In this section, we present in detail the approach we took to numerically solve the Euler-Bernoulli equation in the general scenario of arbitrary deformation magnitudes where the small angle approximation applied in the previous section may no longer be valid. The approach detailed in this section as well as the following one is relevant to Figs. 2-5 in the main text.

Since in general k-fiber profiles may have more than one $y$-intercept at a given $x$ position, we consider a parameterization of the k-fiber shape via an arc length parameter $s$ and solve for $(x(s), y(s))$ instead. Assigning a tangential angle $\theta(s)$ to each point on the k-fiber, we write the Euler-Bernoulli equation as

$$\kappa(s) = -\frac{\mathrm{d}\theta}{\mathrm{d}s} = \frac{M(s)}{EI}. \tag{17}$$

As we can see, knowing the bending moment $M(s)$ at a given arc length position $s$ gives us the rate of change in the tangential angle $\theta(s)$.

To numerically solve for the k-fiber profile, we need initial conditions and update rules. For convenience, we always initialize the first k-fiber point at the origin and therefore set $x(0) = y(0) = 0$. Euler-Bernoulli equation provides us with the derivative of the tangential angle $\theta(s)$. The initial angle $\theta(0)$ therefore needs to be initialized also. When fitting the model to experimental data, we make an educated guess for $\theta(0)$ based on the initial tangential angle of the data profile. Over the course of model optimization, this initial angle is treated as a parameter and is optimized over for better model fitting.

The k-fiber shape profile is solved iteratively using a finite difference method. Specifically, the bending moment $M(s)$ is first used to estimate the tangential angle at position $s + \Delta s/2$ via

$$\theta(s + \Delta s/2) = \theta(s) - \frac{M(s)}{EI}\frac{\Delta s}{2}. \tag{18}$$

Then, the new coordinate on the k-fiber profile is calculated using this angle via

$$x(s + \Delta s) = x(s) + \Delta s\, \cos(\theta(s + \Delta s/2)), \tag{19}$$

$$y(s + \Delta s) = y(s) + \Delta s\, \sin(\theta(s + \Delta s/2)). \tag{20}$$

Finally, the tangential angle is updated for the next step using the bending moment at $s + \Delta s/2$, namely

$$\theta(s + \Delta s) = \theta(s) - \frac{M(s + \Delta s/2)}{EI}\Delta s. \tag{21}$$

Our approach of dividing each step into two half-steps reduces the error in shape calculation, making it quadratic in the step size, i.e., $O(\Delta s^2)$.

For each separate model considered in our work (Fig. 2a, Fig. 3b,d, Fig. 4b, and Fig. 5a), the corresponding expression for the bending moment is used when evaluating $M(s) = M(\vec{r}(s))$. Moment contribution from the microneedle force $\vec{F}_{\mathrm{ext}}$ is considered only when the current position $x(s)$ exceeds the position of the external force $x_{\mathrm{ext}}$. A similar treatment is used also for the point crosslinking force $\vec{F}_c$ applied at position $x_c$. Lastly, in the case of distributed crosslinking shown in Fig. 4b, we account for the integrated moment contribution if the current position $x(s)$ falls in the crosslinking region $(L - \sigma, L)$. Mechanical moments of distributed crosslinking forces up to position $x(s)$ are calculated by treating them as a series of linear Hookean springs exerting a restoring force $-ky(s')\Delta x'$ on k-fiber segments $(s', s' + \Delta s)$ with an $x$-projection size $\Delta x'$. Here, $k$ is the effective 'spring constant' chosen in our studies to be sufficiently large to result in a negative curvature response near the kinetochore.

### D Model fitting procedure

We obtain the best model fits to the extracted k-fiber profiles by minimizing the sum of squared errors. Error is defined for each data point of the tracked k-fiber as the minimal distance between that point and the model k-fiber profile, with the model profile represented as a piecewise linear curve (Fig. S5). This error metric can be successfully applied to profiles with sophisticated shapes, as opposed to the more traditional metrics based on errors in $y(x)$ prediction which become ill-defined for curved profiles with more than one $y$-value at a given $x$-position.

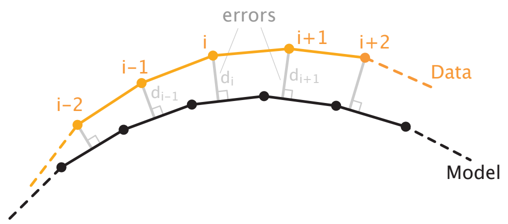


Figure S5: Schematic of the error definition. For the $i^{\mathrm{th}}$ data point, the error $d_i$ is the smallest distance to the model profile which is represented as a piecewise linear curve. Sum of squared errors, namely, $\sum_i d_i^2$, is minimized in the model fitting procedure.

During parameter search, we impose constraints on the parameter values and possible k-fiber shape profiles based on our understanding of the experimental setup. This helps us avoid physically unrealistic scenarios. Below we list the main constraints that we imposed:

- The inferred microneedle force has to point outward, with its $y$-component having the same sign as k-fiber deflection, i.e., it has to be positive.
- The inferred microneedle force is perpendicular to the local tangent of the k-fiber shape profile. This is based on our assumption of very low frictional forces in the tangent direction discussed in the Materials and Methods.
- The inferred position of microneedle force application is within 0.5 µm of the position of profile peak. We give this finite range in our search for the effective point of force application because the precise point of contact between the k-fiber and the microneedle is hard to identify from fluorescence microscopy images, and because the contact likely takes place over a small but finite contour length.
- If the microneedle force has a positive $x$-component (points to the right), then the point force at the left k-fiber endpoint ($F_x$) has to have a negative $x$-component (point to the left) in order to balance the external force. This condition is imposed to avoid considering spontaneous inward-pointing buckling forces during parameter search. An analogous constraint is applied if the microneedle force has a negative $x$-component (points to the left).
- Parameter sets that predict k-fiber shape profiles with loops (i.e., the model curve passes through the same point twice) are not considered during search.

Because of this set of hard constraints, standard gradient descent-based methods for minimizing the sum of squared errors are not effective for finding the optimal parameters. We therefore use a time consuming but more reliable stochastic search method. There, we initialize multiple "walkers" at different positions in the parameter space, and run a stochastic search with up to 150,000 steps where each step is more likely to be taken in the direction that decreases the overall error. The best model fit is then generated by the set of sampled parameters that yields the lowest sum of squared errors.

### E Modeling of PRC1 binding states

In this section, we provide the details of our approach to distinguish the PRC1 populations by their binding state using equilibrium thermodynamic modeling and immunofluorescence imaging data (Suresh et al., 2020). The results presented here are relevant to Fig. 4i of the main text.

We distinguish three binding states for PRC1 - freely diffusing, singly bound, and doubly bound. The freely diffusing population represents the unbound PRC1 molecules that occupy the entire volume of the cell and do not contribute to crosslinking activity. We denote the concentration of this PRC1 population by $c_f$ and assign no spatial dependence to it, since intracellular diffusion occurs at much faster time scales than metaphase and would therefore manage to equilibrate the free PRC1 population in the cell.

The singly bound population includes PRC1 molecules that are bound to a single microtubule only and, similar to the free population, do not contribute to crosslinking activity. We denote this population by $c_1(\vec{r})$ and relate it to the local tubulin concentration $\rho_{\mathrm{MT}}(\vec{r})$ via

$$c_1(\vec{r}) = \frac{\rho_{\mathrm{MT}}(\vec{r})}{K_d} c_f, \tag{22}$$

where $K_d$ is the dissociation constant of PRC1–single microtubule binding. To write the above relation between the free and singly bound populations, we again considered an equilibrated scenario, which we assume holds true given the fast dynamics of molecular turnover (Pamula et al., 2019) and diffusion compared to the duration of metaphase.

If $c_{\mathrm{tot}}(\vec{r})$ is the local concentration of all PRC1 populations together, then the doubly bound PRC1 population ($c_2(\vec{r})$) can be isolated by subtracting the free and singly bound populations from the total one, namely

$$\begin{aligned} c_2(\vec{r}) &= c_{\mathrm{tot}}(\vec{r}) - c_1(\vec{r}) - c_f \\ &= c_{\mathrm{tot}}(\vec{r}) - \frac{\rho_{\mathrm{MT}}(\vec{r})}{K_d} c_f - c_f. \end{aligned} \tag{23}$$

We are interested in estimating $c_2(\vec{r})$ along the pole-pole axis of the spindle in order to infer the length scale of the active crosslinking region.

To that end, for each spindle, we first estimate $c_f$ by averaging over the measured immunofluorescence in several different regions of interest (ROIs) where there is little to no detectable presence of microtubules. Examples of such ROIs are shown in Fig. S6a. Next we need to estimate the dissociation constant $K_d$. Based on the *in vitro* measured ≈30-fold higher binding affinity of PRC1 to antiparallel microtubules compared to parallel ones (Bieling et al., 2010), and the result of an electron microscopy study suggesting that microtubules near the spindle poles are predominantly parallel (Euteneuer and McIntosh 1981), we assume that the PRC1 population in the immediate vicinity of spindles poles is made out of free and singly bound states only. Denoting the pole-proximal positions by $\vec{r}_p$, we set $c_2(\vec{r}_p) \approx 0$ and use Eq. 23 to estimate $K_d$ as

$$K_d \approx \left\langle \frac{\rho_{\mathrm{MT}}(\vec{r}_p)\, c_f}{c_{\mathrm{tot}}(\vec{r}_p) - c_f} \right\rangle_{\vec{r}_p}, \tag{24}$$

where $\langle \cdot \rangle$ represents averaging over pole-proximal positions $\vec{r}_p$. Manually selecting several ROIs near the poles (Fig. S6b) and using the immunofluorescence measurements for $c_{\mathrm{tot}}(\vec{r}_p)$ and $\rho_{\mathrm{MT}}(\vec{r}_p)$ in these regions, we perform the averaging and obtain the estimate for $K_d$.

With $c_f$ and $K_d$ calculated, we obtain the spatial profiles of actively engaged PRC1 molecules along the pole-pole axis of the spindle by selecting a rectangular region spanning the area between the poles (Fig. S6c) and using the measured PRC1 ($c_{\mathrm{tot}}(\vec{r})$) and tubulin ($\rho_{\mathrm{MT}}(\vec{r})$) profiles to calculate $c_2(\vec{r})$ via Eq. 23. Lastly, approximating k-fibers as homogeneous bundles of microtubules, we divide the calculated concentration of actively engaged PRC1 molecules by the local tubulin concentration, and report that ratio (engaged PRC1 per tubulin – a proxy for the strength of local crosslinking) as a function of position in the main text (Fig. 4i).

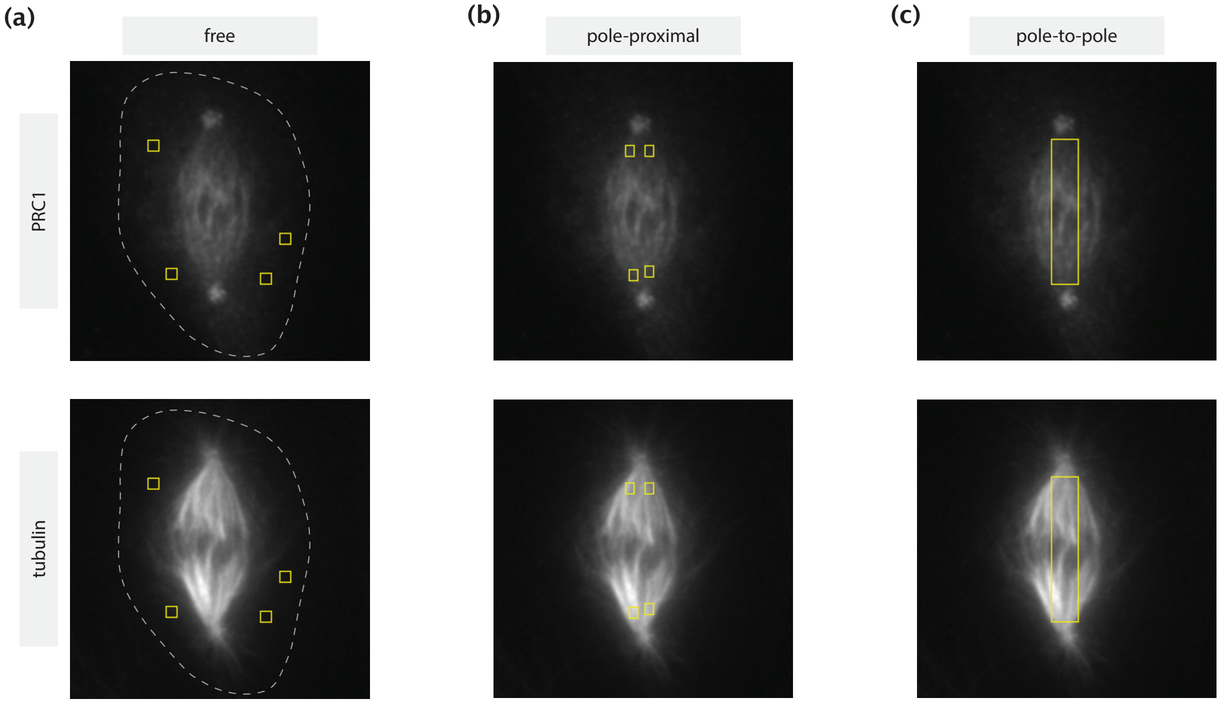


Figure S6: ROIs selected for different calculations shown on immunofluorescence images of PRC1 (top row) and tubulin (bottom row). (a) Regions with little to no tubulin presence where PRC1 can be considered unbound. The dashed lines represent the cell boundaries estimated by manual tracing based on high intensity contrast. (b) Pole-proximal regions where microtubules are present primarily in a parallel configuration. (c) Rectangular pole-to-pole region where the estimation of the actively engaged PRC1 population is made.
